# Broad misappropriation of developmental splicing profile by cancer in multiple organs

**DOI:** 10.1101/2021.12.13.472504

**Authors:** Arashdeep Singh, Arati Rajeevan, Vishaka Gopalan, Piyush Agrawal, Chi-Ping Day, Sridhar Hannenhalli

**Affiliations:** Cancer Data Science Lab, National Cancer Institute, National Institutes of Health; Laboratory of Cancer Biology and Genetics, National Cancer Institute, National Institutes of Health

## Abstract

Oncogenesis mimics key aspects of embryonic development. However, the underlying molecular determinants are not completely understood. Leveraging temporal transcriptomic data during development in multiple human organs, we demonstrate that the ‘embryonic positive (EP)’ alternative splicing events, specifically active during human organogenesis, are broadly reactivated in the organ-specific tumor. EP events are associated with key oncogenic processes and their reactivation predicts proliferation rates in cancer cell lines as well as patient survival. EP exons are significantly enriched for nitrosylation and transmembrane domains coordinately regulating splicing in multiple genes involved in intracellular transport and N-linked glycosylation respectively, known critical players in cancer. We infer critical splicing factors (CSF) potentially regulating these EP events and show that CSFs exhibit copy number amplifications in cancer and are upregulated specifically in malignant cells in the tumor microenvironment. Mutational inactivation of CSFs results in decreased EP splicing, further supporting their causal role. Multiple complementary analyses point to MYC and FOXM1 as potential transcriptional regulators of CSFs in brain and liver, which can be potentially targeted using FDA approved drugs. Our study provides the first comprehensive demonstration of a splicing-mediated link between development and cancer, and suggest novel targets including splicing events, splicing factors, and transcription factors.

## Introduction

Cancer onset and progression results in the dedifferentiation and gradual loss of lineage-specific phenotypes and echoes multiple facets of early embryonic development including rapid proliferation, epithelial-mesenchymal transition (EMT), cellular migration, and angiogenesis. The mechanistic details of these cancer-associated changes in cellular function and physiology, termed as ‘hallmarks of cancer’ (Hanahan & Weinberg, 2011), are not completely understood. Past studies have shown that a core set of transcription factors (TFs) and signaling pathways, which maintain pluripotency in embryonic stem cells (ESCs) and orchestrate normal embryonic development, are reactivated in cancer and thus underlie physiological reversal in cancer progression (Ben-Porath et al., 2008; Kelleher et al., 2006; Kim et al., 2010; Patra, 2020). For instance, the core pluripotency markers Oct3/4 and Sox2, are important biomarkers of several cancers (Gidekel et al., 2003; Li et al., 2004; Rodriguez-Pinilla et al., 2007). Likewise, the Myc module of ESCs gets reactivated in mouse models of mixed-lineage leukemias and is a predictor of patient outcome in many human cancers (Kim et al., 2010). Consistent with these anecdotes, a universal signature of stemness accurately predicts the tumor infiltration by leukocytes and response to immunotherapy(Malta et al., 2018). In addition to TFs, various signaling pathways involved in embryonic development, such as Wnt, Notch, Hippo, also get reactivated in cancer and their associated genes accumulate oncogenic mutations (Dreesen & Brivanlou, 2007; Kelleher et al., 2006; Sanchez-Vega et al., 2018).

In addition to gene expression, alternative splicing (AS), wherein multiple isoforms of the same gene are expressed, affects >95% of the multi-exonic genes in humans (Pan et al., 2008; E. T. Wang et al., 2008) and underlies diverse biological processes such as stemness, differentiation, development, and ageing (Agosto & Lynch, 2018; Baralle & Giudice, 2017; Han et al., 2013; Salomonis et al., 2009). A plethora of gene-centric studies have demonstrated the critical role that AS plays in cancer (Bonnal et al., 2020). For instance, long and short isoforms of Bcl-x protein have anti-apoptotic and pro-apoptotic roles respectively (Takehara et al., 2001; Xerri et al., 1996). Several members of the receptor tyrosine kinase family express multiple isoforms enhancing the proliferative or metastatic ability of cancer cells. For example, the FGFR2 isoform, FGFR2III-b, is mainly expressed in epithelial cells while FGFR2III-c is expressed in mesenchymal cells (Yan et al., 1993). This isoform switching is involved in epithelial-mesenchymal transition (EMT) (Warzecha et al., 2009) and is linked to invasiveness and metastasis of colorectal (Matsuda et al., 2012; Sonvilla et al., 2010) and breast cancers (Cha et al., 2008). Likewise, alternatively spliced isoforms of genes such as P63, Cyclin D1, CD44, H-Ras, Rac1, PKM etc. can modulate proliferative, apoptotic, metabolic, and invasive properties of cancer cells (Bonnal et al., 2020; David & Manley, 2010; Pokorná et al., 2021).

Recent comparative transcriptomic analyses across multiple organs showed the prevalence and cross-species conservation of alternative splicing events during development (Mazin et al., 2021). Despite the established importance of AS in development and cancer, as well as broad phenomenological links between development and cancer, an unbiased and comprehensive investigation of the links between development and cancer AS events in a tissue-specific fashion is still lacking and can have major implications on our broader mechanistic understanding of oncogenesis and cancer therapies. Leveraging the human developmental transcriptome across multiple time points in three organs (Cardoso-Moreira et al., 2019) as well as the transcriptomic data of the corresponding cancer from TCGA (https://www.cancer.gov/tcga), here we chart the landscape of embryonic splicing events that are reactivated in the organ-specific cancer, and investigate their upstream regulators and downstream functional implications. Focusing on the most common type of AS event type, namely, exon skip events, we show that embryonic AS events associate with key oncogenic processes such as rapid proliferation, migration, and angiogenesis, and are significantly reactivated in tumors. The reactivation of embryonic AS events predicts the patient’s survival and is associated with the proliferation rate in CCLE cell lines. We found that nitrosylation domain (ND), transmembrane-region domain (ND), and WD40 domain are significantly enriched among EP and EN exons in all three tissues. Detailed molecular and functional analysis reveals that NDs and TRDs respectively affect retrograde cellular transport by coordinately regulating the activity of Arf and Ras family GTPases and N-linked glycosylation by regulating the transmembrane localization of oligosaccharyl transferase subunits.

We further trained a splicing regulatory model based on the developmental data which accurately predicts the inclusion of embryonic AS events in cancer patients and identifies critical splicing factors (CSFs) potentially regulating embryonic AS events. We find that identified CSFs are upregulated in cancer, often accompanied by copy number amplifications. Based on multiple complementary approaches, we also identify key transcription factors (TFs) predicted to regulate the identified SFs, including MYC, FOXM1 in the brain and liver, respectively. Leveraging tumor scRNA-seq data, we show that the CSFs are specifically activated in the malignant epithelial cells, further supporting their role in malignancy. Finally, through drug repositioning approaches, we provide a list of FDA approved drugs potentially targeting the TF regulators of CSF genes. Overall, our work firmly establishes, through multi-modal data integration, reversal to developmental AS in cancer, and suggests novel therapeutic avenues directly targeting the regulators of such a reversal.

## Results

### Identification of exons associated with human fetal development

To identify the AS events associated with fetal development, we implemented a two-step approach where we first identified fetal-associated pathways, and then obtained the AS events correlated with those pathways (Fig. 1A); the rationale and advantages of this approach are discussed in the Methods section. Based on organ-specific transcriptomic data across multiple stages (Supplementary table S1) of pre- and post-natal development (Cardoso-Moreira et al., 2019), we first estimated the activity for each of the 332 KEGG pathways (Kanehisa et al., 2012), quantified as the median expression of the pathway genes, in each sample, independently in brain, liver and kidney tissues. Principal component analysis (PCA) of the pathway activity clearly separates the pre- and post-natal stages along the first principal component (Supplementary Fig. S1A). Clustering of pathways in the PCA space (Methods) revealed two mutually exclusive sets of pre- or post-natal pathways which were correspondingly assigned as ‘embryonic positive’ or ‘embryonic negative’ (Fig. 1B).

**Figure 1.**
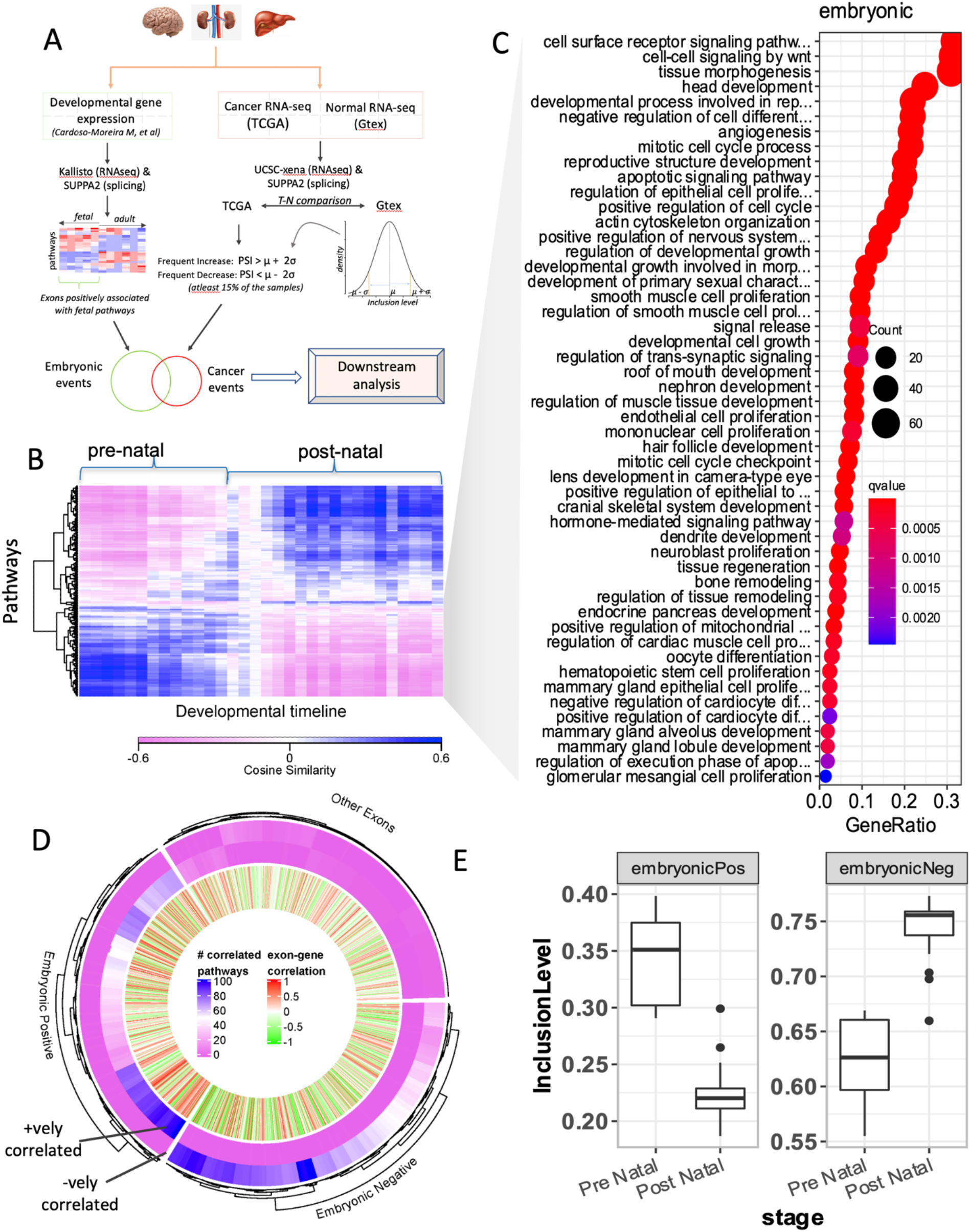
Detection of AS events relevant to development of organs. (A) Overview of the pipeline for the identification and comparison of developmental and cancer-associated splicing. (B) Hierarchical clustering of KEGG pathways in brain cerebellum. (C) GO enrichment of the genes comprising embryonic pathways in (B). (D) Circular heatmap showing the number of positively and negatively correlated embryonic pathways with each exon. Each leaf in the dendrogram is an exon. Outer two rows represent respectively the number (per legend colors) of positively and negatively correlated embryonic pathways with the exon. The innermost layer shows the correlation of the PSI value of each exon with its cognate gene. For visual clarity, only 1000 randomly chosen exons are included in the plot. (E) Boxplots showing the differential inclusion of embryonic positive (EP) and embryonic negative (EN) events during pre- and post-natal stages of development.

As expected, genes constituting embryonic positive pathways are enriched in several gene ontology (GO) terms related to the well-established oncogenic processes such as EMT, extracellular matrix (ECM) remodeling, cellular proliferation, and angiogenesis, validating the role of embryonic positive KEGG pathways in embryogenesis and organ development (Fig. 1C, Supplementary Table S2). Next, we used PEGASAS (Phillips et al., 2020) to identify alternative exons whose sample-specific inclusion is significantly correlated with the activity of embryonic positive pathways across developmental timepoints (Methods). We defined an exon as embryonic positive (EP) or embryonic negative (EN) based on the fraction of embryonic positive pathways whose activities are respectively significantly positively or negatively correlated with the exon’s inclusion level (Fig. 1D, E, Methods). We thus identified on average ~2000 EP as well as EN exon skip events in each tissue (Supplementary Table S3); as expected, EP and EN exons exhibit mutually exclusive inclusion patterns in the pre- and postnatal stages (Fig. 1E, and Supplementary Fig. S1B).

We found that the EP and the EN exon inclusion levels are broadly uncorrelated with the expression of their cognate genes, suggesting that these AS events are independent of their cognate gene’s expression (Fig. 1D, Supplementary Fig S1C). This independence is further supported by our observation that ~20-30% of the cognate genes of EP/EN exons in fact contain both EP and EN events (Supplementary Fig. S1D). Moreover, in almost all cases (> 99%) when an exon’s inclusion correlates with an embryonic positive pathway, the exon’s cognate gene is not member of that pathway. Collectively, these data suggest that AS provides an additional regulatory layer to gene expression programs for controlling developmental pathways.

The cognate genes of EP and EN exons are significantly enriched in tissue specific processes in the case of brain and liver (Supplementary Fig. S1E, Supplementary Table S4). For example, GO terms for neuronal activities, such as synapse organization, dendrite development, neuron death, cell polarity, regulation of neurotransmitters, are enriched in the cognate genes of EP/EN exons specifically in brain. Likewise, liver EP/EN exons are involved in the regulation of many key metabolic processes as well as regulation of cell junctions and cytokinesis. EP/EN exons in all three tissues are enriched for autophagy, consistent with the emerging role of AS in the regulation of autophagy (Paronetto et al., 2016). Overall, we identify numerous exons that, independent of the expression of their cognate gene, are preferentially utilized during fetal development and repressed postnatally, and strongly associate with key developmental and oncogenic processes.

### Embryonic AS events are recapitulated in cancer and are associated with cancer stage and patient survival

We next assessed the extent to which the organ-specific EP events are recapitulated in the corresponding cancer types. First, we found that in all three organs the genome-wide profile of the AS events clearly distinguishes tumor samples from their non-malignant counterparts in GTEx (Supplementary Fig. S2A), as observed previously (Phillips et al., 2020; Tsai et al., 2015). Next, we identified the cancer-associated AS events in each organ by comparing the splicing profiles in tumors with healthy GTEx counterparts (Methods, Fig. 1A) and assessed their overlap with organ-specific EP and EN events. In all three organs we found that the EP events are significantly enriched among the AS events frequently increased in cancer, while the EN events are enriched among the AS events frequently decreased in cancer (Fig. 2A). The observed enrichment may simply be because EP events have lower inclusion level in healthy postnatal tissues as shown in Fig. 1E, and are therefore more likely to increase in cancer (analogously EN events might be more likely to decrease). We ruled out this potential confounder by randomly sampling exons with low (psi < 0.3) and high (psi >0.7) inclusion level in healthy GTEx samples of liver and testing their enrichment among events frequently increased and decreased in liver cancer, respectively (nominal false positive rate < 0.01; Supplementary Fig. S2B; Methods). Interestingly, we observe an even greater enrichment of EP and EN events in advanced tumors compared with early-stage tumors (Methods; Supplementary Fig. S2C,D), strongly linking embryonic splicing to not only oncogenesis but also to cancer progression. Furthermore, in all three organs, the EP (respectively EN) inclusion levels across samples are positively (respectively, negatively) correlated with cancer hallmark signature gene set scores (Fig. 2B), indicating a possible direct link between oncogenic processes and embryonic splicing. Unlike other signatures, apoptosis and DNA damage gene sets, whose activity is known to inversely correlate with tumor aggressiveness (Hanahan & Weinberg, 2011), are negatively correlated with EP events.

**Figure 2.**
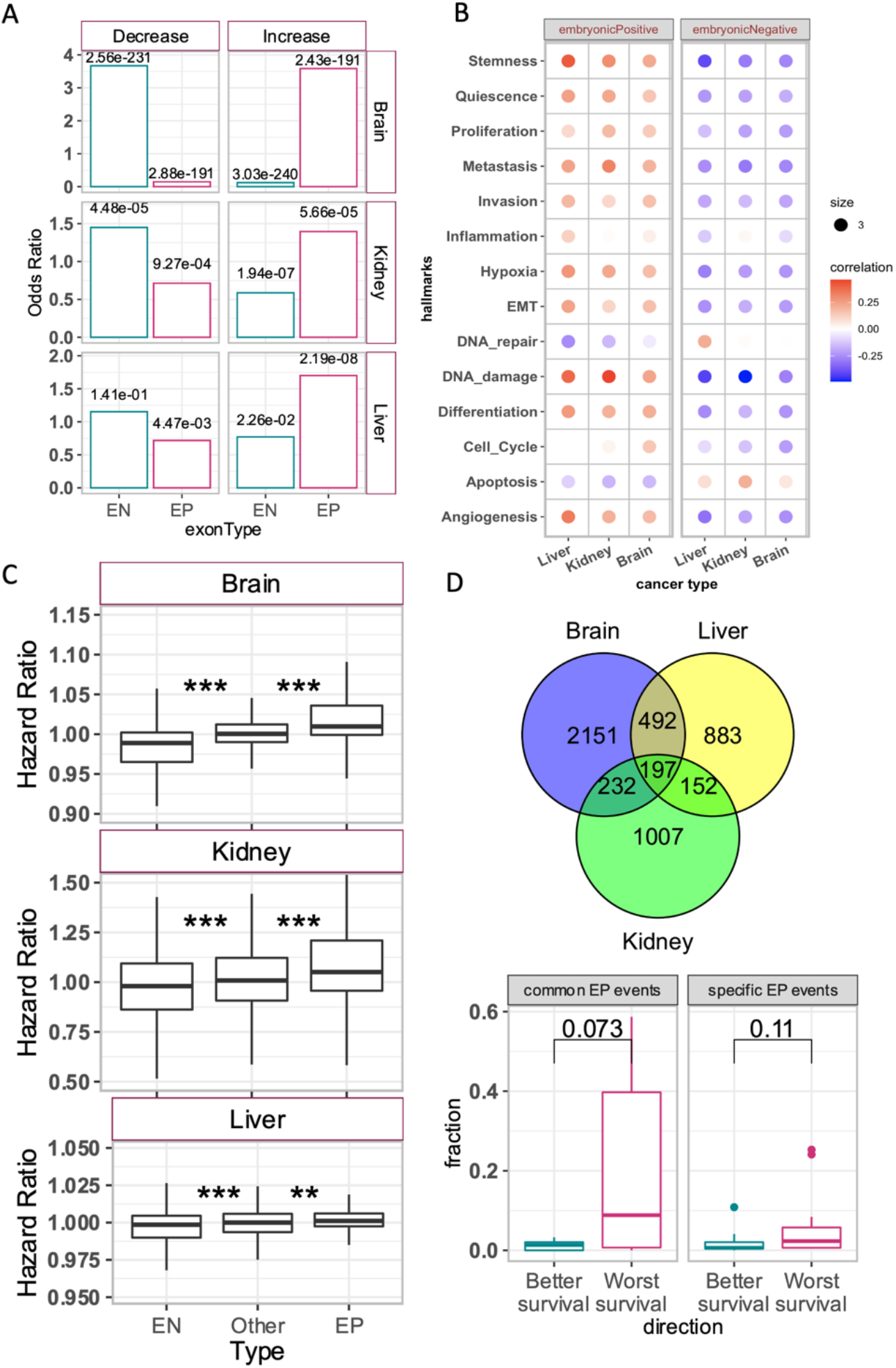
Embryonic splicing events in cancer. (A) Barplots showing the odds ratio of overlap between embryonic splicing events and frequently increased/decreased events in cancer for brain, kidney and liver. The numbers at the top of each bar are FDR corrected values from Fisher test. (B) Dotplots for correlation between the median inclusion level of EP and EN events and cancerSEA hallmark gene sets. (C) Boxplots distribution of hazard ratios of EP and EN detected in brain, kidney and liver in their corresponding cancers. The ‘Other’ set of exons are the remining exon and serve as genome wide control. (D) Venn diagram shows the overlap between the EP events detected in three tissues. (E) Boxplots showing the proportion of EP events associated with better or worst survival among the common (detected in all three tissues) and specific events (detected in only 1 tissue). P-values are from Wilcoxon test.

Next, we directly assessed whether the EP and EN inclusion level is associated with patient survival using Cox regression (Methods). In all three tissues, EP inclusion had significantly higher (and positive) hazard ratios and EN inclusion had respectively lower (and negative) hazard ratios compared to the rest of the exons (Fig. 2C); we ensured that the observed trends were not confounded by the expression level of the cognate genes (Supplementary Fig. S2E).

Since early embryonic development shares several molecular programs across organs (Cardoso-Moreira et al., 2019), we derived a set of EP events (197 events) common to all three organs and assessed its association with survival across 20 cancer types. We found that the shared EP events were a significantly stronger predictor of pan-cancer patient survival as compared to tissue-specific EP events (Fig. 2D), further underscoring the embryonic roots of splicing changes in cancer.

### Alternatively spliced transmembrane-region and nitrosylation domain may regulate N-linked glycosylation and retrograde cellular transport during development and cancer

To get insights into the functions potentially affected by dynamic inclusion of EP and EN exons, we performed molecular functional enrichment analysis of the genes containing the EP and EN events. In all organs, we observed a significant enrichment of Ras GTPase binding, cell adhesion, and cytoskeleton binding classes such as cadherin, actin, and microtubules (Supplementary Fig. S3A). Brain and Liver EP/EN genes were additionally enriched for dynactin and clathrin binding (Supplementary Fig. S3A). These processes promote tumorigenesis by modulating the cytoskeleton and cellular transport during the proliferation and migration of cancer cells (Boudhraa et al., 2020; Brayford et al., 2015). A more detailed discussion is provided below in the Discussion section.

To gain further insights into the molecular role of EP and EN exons and investigate their link with oncogenesis, we identified protein domains from PFAM database (El-Gebali et al., 2019) enriched among the EP/EN exons (Methods). Three domains – transmembrane-region domain (TRD), nitrosylation domain (ND), and WD40 – are enriched among EP and EN exons in all three organs (Fig. 3A), leading us to speculate their potential role in some of the functions performed by the cognate genes of EP and EN exons. To explore this potential link, we identified the gene subsets whose EP/EN exons contained these domains (total 6 gene subsets per tissue: 3 domains x 2 EP/EN gene sets) and performed molecular function enrichment analysis for each subset (Figure 3B). As expected, enriched molecular functions in a gene set could be unambiguously attributed to the corresponding domain. For instance, gene subsets of WD40 domains were enriched for ubiquitin binding, consistent with the established role of WD40 as binding interfaces for ubiquitin proteins (Pashkova et al., 2010). Likewise, the genes containing the transmembrane region domain were indeed enriched for various kinds of transmembrane transporters (Fig. 3B). Further, the assessment of overlap among the cognate genes of EP and EN exons harboring these domains across tissues indicates that observed enrichment of protein domains is not driven by the same set of genes but instead, multiple genes coordinate the splicing of these domain across tissues (Fig S3B). To probe the interplay between these enriched molecular functions and biological processes affected by dynamic inclusion of these domains, we performed biological processes enrichment analysis on the same gene sets and assessed the overlap of genes having a specific enriched molecular function with those having a specific enriched biological process.

**Figure 3.**
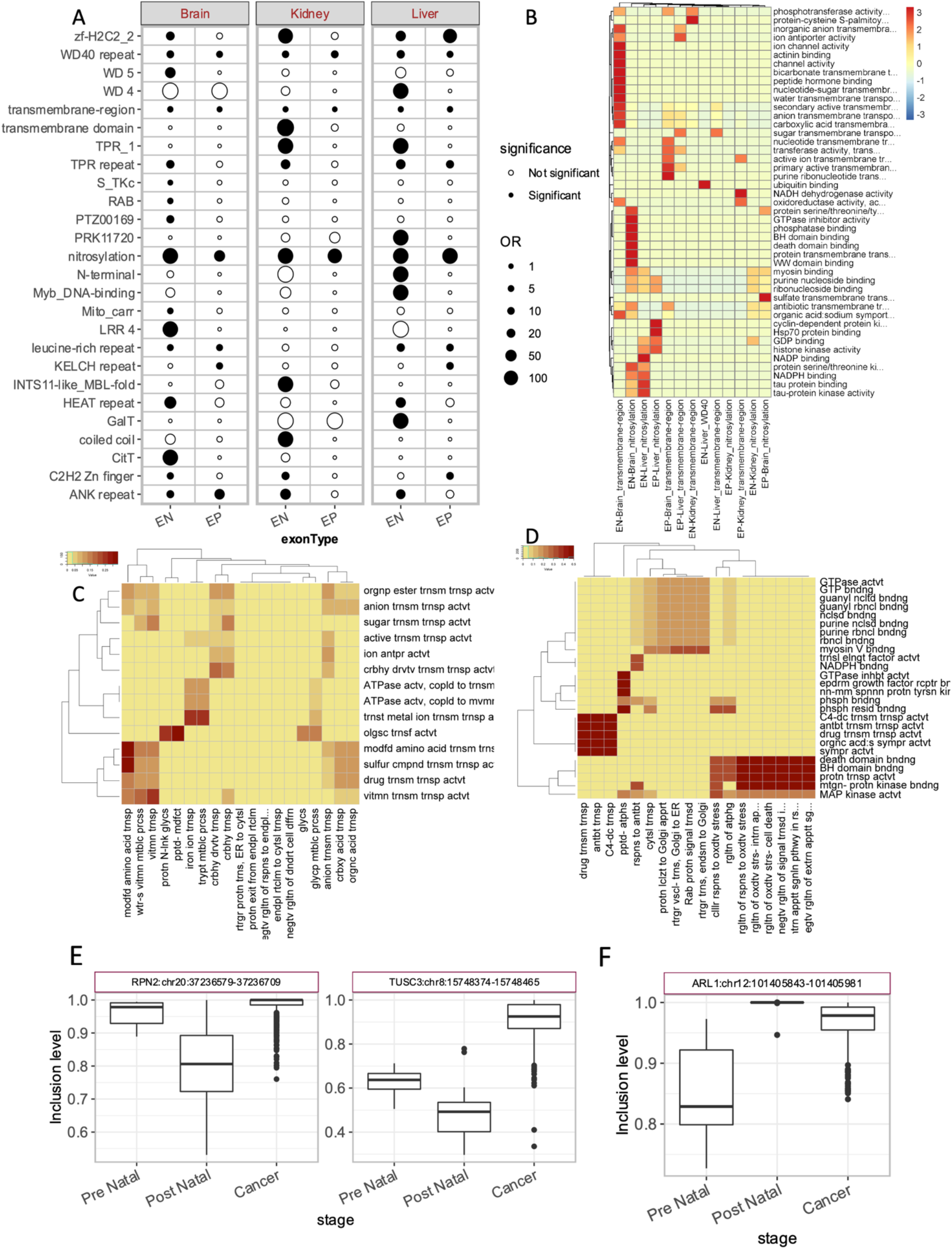
Functional assessment of EP and EN exons. (A) Dotplot showing the enrichment of domains in EP and EN events across three tissues. Size of dots is scaled according to the magnitude of odds ratio and solid and hollow dots respectively indicate significant and non-significant domains. (B) Molecular functional enrichment across three organs for the cognate genes of EP and EN events containing nitrosylation, transmembrane-region and WD40 domains (indicated along the columns). The heat colors indicate −log10 of adjusted p-value. (C) Heatmap showing the cooccurrence of enriched biological process (columns) and molecular functions (rows) among the genes containing transmembrane-region domain in EP exons in liver. (D) same as (C) but for nitrosylation domain in EN exons in brain. (E &F) The inclusion of EP exons with TRD among the subunits of OST complex (E) and EN exons with ND among the GTPases potentially involved in the regulation of vesicle transport (F)

The observed correspondence between molecular function and biological processes among the cognate genes of EP and EN exons is well supported. For instance, in brain, cognate genes of EN exons with a transmembrane domain and encoding various types of transporters (molecular function) are predominantly involved in cross-membrane transport (biological process) (Fig. 3C).

Moreover, in brain and liver EP exons, the molecular function oligosaccharyl transferase activity significantly overlapped with biological processes related to N-glycosylation of proteins (Fig. 3C, and S3C), a modification which typically takes place in the phospholipid bilayer of ER and Golgi bodies through the multi-subunit oligosaccharyl transferase complex (OST). Remarkably, four subunits of OST showed a coordinated reduction in the inclusion of TRD from pre- to post-natal stages, which increased again in cancer patients in brain (Fig. 3E), with TUSC3 and RPN2 undergoing greatest change. This suggests that modulation of transmembrane localization of OST through alternative splicing of TRD during embryogenesis might directly impact the process of N-glycosylation. Interestingly, N-glycosylation of several proteins have been implicated in cellular proliferation and migration by modulating the cell-matrix interactions (Pinho & Reis, 2015). Therefore, increased inclusion of TRD among the subunits of OST might help the cancers (Fig. 3E) to upregulate the increased demand for N-glycosylation.

Similar analysis for ND revealed that cognate genes of EN exons containing this domain in the brain were significantly enriched for the molecular functions related to GTPase activity and its regulators (Fig. 3B). Previous studies have strongly implicated nitrosylation in the upregulation of GTPase activity (Lin et al., 2018; Raines et al., 2007). Our result thus suggests a novel role of alternatively spliced ND in the modulation GTPase activity during embryogenesis, and cancer. In fact, few of the genes containing nitrosylation domain among brain EN exons, such as RAB6A and RAB6B, are themselves GTPases belonging to RAS oncogene family, hinting at autoregulation of their GTPase activity through dynamic inclusion and exclusion of nitrosylation domain. As for transmembrane domain, we obtained the genes having a ND among the EN exons in brain and identified the correspondence between the enriched molecular functions and biological processes (Fig. 3D). We observed that genes having GTPase activity were involved in Rab protein signal transduction and retrograde vesicle transport from endosomes to Golgi bodies to endoplasmic reticulum (Fig. 3D), the processes where GTPases are known to play a critical role (Chi et al., 2013; Suda et al., 2018). Among the GTPases having a ND in their EN exons, ARL1 gene had the largest change in the inclusion of ND from pre-natal to post-natal stages and then in cancer (Fig 3F). Our analysis thus suggests the novel role of alternatively spliced ND in the regulation of the cytoplasmic transport by modulating the activity of GTPases. Additionally, some of the genes containing a ND among brain EN exons were enriched for the molecular functions related to BH-domain binding, death-domain binding and MAP-kinase signaling, which corresponded to the processes related to intrinsic apoptotic signaling pathways (Fig. 3D), potentially implicating exclusion of nitrosylation domain in modulating apoptosis (Iyer et al., 2008). Overall, our results implicate recapitulation of embryonic alternative splicing patterns of transmembrane and nitrosylation domains in several key oncogenic processes.

### Splicing regulatory model of EP events reveals key splicing factors dysregulated in cancer

Splicing factors (SF) control the choice and inclusion level of exons (Blanchette et al., 2005). To identify potential SFs regulating embryonic splicing, we trained a partial least squares regression (PLSR) model to predict the overall inclusion level of EP events based on the expression levels of 442 annotated SFs (Methods, Supplementary Table S5). In each organ, trained solely on the developmental data, our model predicted the median inclusion level of EP events in independent tumor samples with remarkably high accuracy (average correlation between predicted and observed EP levels ~0.88; Fig. 4A).

**Figure 4.**
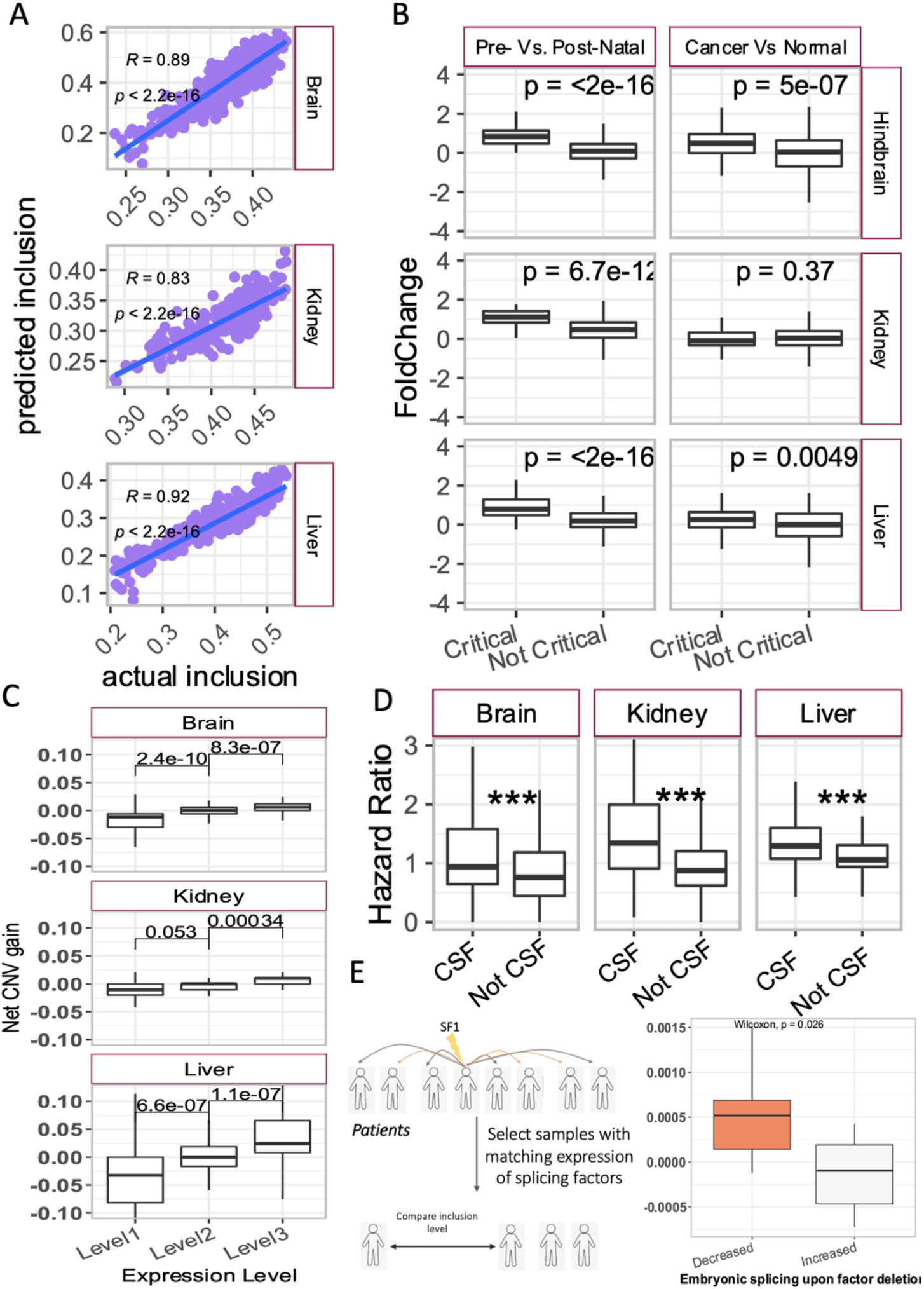
Splicing regulatory model of EP events reveals key splicing factors dysregulated in cancer. (A) Scatter plot of the actual and predicted median inclusion level of EP events across TCGA samples in a tissue-specific cohort. The numbers in each plot area denote the Pearson correlation and the p-value. (B) Boxplot distribution of fold-change of critical and non-critical splicing regulators of EP splicing events during development (left) and in cancer (right). (C) Boxplots showing distributions of net CNVs in critical splicing factors in patients stratified based on the expression of the splicing factors. (D). Boxplots showing the distributions of Cox hazard ratios computed using the gene expression of critical and control splicing factors potentially regulating EP. (E) Schematic illustration of mutation analysis (left) and the distribution of the regression coefficients (y-axis) of splicing factors resulting in decrease or increase in the median inclusion level of EP events in the mutated samples compared to expression matched unmutated samples. In all boxplots, p-values are from Wilcoxon test.

As expected, the top positive predictors of EP events identified based on their coefficients in the PLSR model, which we refer to as critical splicing factors (CSFs, Supplementary Table S5), had higher expression during the prenatal stage of development, and underwent significant upregulation in their corresponding cancer (Fig. 4B). Further, the deletion of orthologous genes of brain CSFs result in defective nervous system development in mice, and CSFs from all three tissues are much more likely to result in preweaning lethality as compared to nCSFs (Fig S4A), further supporting the developmental role of CSFs.

Interestingly, cancer patients with higher expression of CSFs and correspondingly higher inclusion level of EP events have a significantly greater number of copy number amplifications in CSFs (Fig. 4C). In addition, a gain in CSF expression is significantly associated with worse patient survival (Fig. 4D).

To assess whether CSFs play a causal role in regulating EP events, we tested if the EP inclusion level is lower in tumor samples bearing nonsense (inactivating) mutations in CSFs. We first identified all SFs whose mutant samples have lower EP inclusion than the wildtype samples and found that such potentially causal SFs have significantly higher (and positive) regression coefficients as compared to the other SFs (Fig. 4E), establishing a potentially causal role of CSFs in regulation of EP events. We ensured that our results are not confounded by SF expression differences between the mutant and wildtype samples (Methods). Overall, these results reveal potentially causal SFs underlying the EP events and link induction of such SFs, potentially via copy number amplification, to cancer.

### Embryonic splicing events are associated with proliferation rates in cancer cell lines

Our results above (Fig. 1B, 2A) suggest that increased inclusion of EP events in tumors might be involved in mediating oncogenic processes such as rapid proliferation, EMT, and angiogenesis. Leveraging the DepMap database (https://depmap.org/portal/) that includes RNA-seq data and proliferation rates in multiple cancer cell lines, we find that in liver and brain, there is a remarkable association between the doubling time and the EP/EN inclusion levels across cell lines derived from the organ-specific cancer type (Fig. 5A).

**Figure 5.**
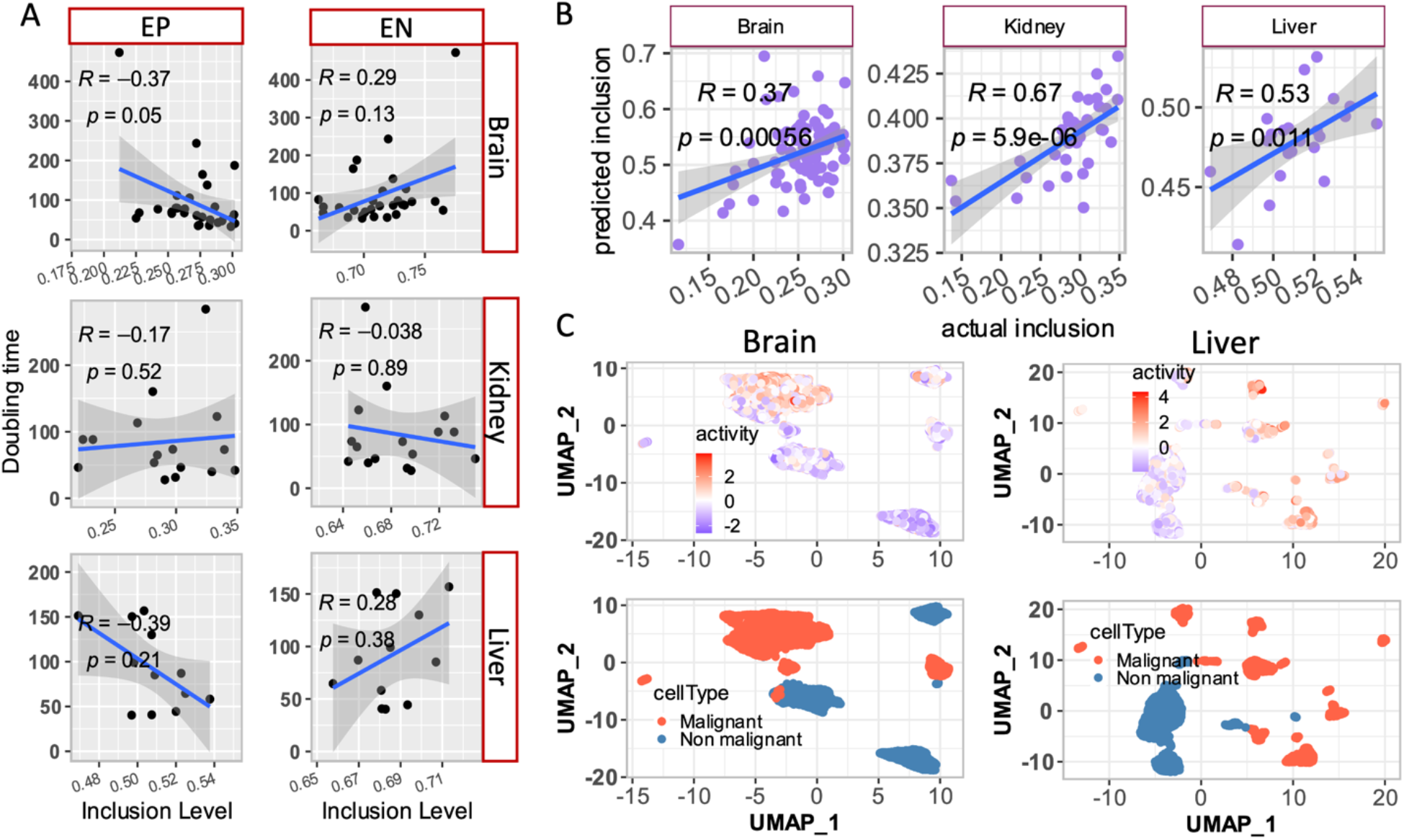
Embryonic splicing events and their regulators in cancer cell lines. (A) Scatter plot of median inclusion level of EP and EN events with the doubling time of cancer cell lines (obtained from Depmap portal) corresponding to brain, kidney, and liver. Pearson correlation and p-value are shown. (B) Scatter plot of the observed and predicted median inclusion level of EP events in brain, kidney and liver cancer cell lines. (C). UMAP showing the activity of critical splicing factors in the malignant and non-malignant cells of the tumor microenvironment; top row: activity, bottom row: cell types.

Further, splicing regulatory models learned from the developmental data accurately predict the EP event inclusion in the corresponding cell lines (Fig. 5B). Collectively, these observations further validate the links between CSFs and proliferation, mediated by EP events. Given the links between CSF activity and proliferation, we expect that inactivation (by CRISPR or RNAi) of the CSFs will have an adverse effect on the proliferation rates of the cell lines. Indeed, we found that in liver cancer-derived cell lines, the more critical a SF (based on PLSR coefficient), the greater was the dependency of the cell line on that SF (negative dependency scores, Supplementary Fig. S5), supporting a functional role for CSFs; however, we did not see this trend in brain and kidney, as discussed below. Further supporting the role of CSFs in malignant transformation, we found that in the single-cell transcriptome of liver and brain (Methods) tumor microenvironment, CSFs were specifically expressed in the malignant but not in non-malignant cells (Fig. 5C). Collectively, these observations link the role of CSFs in tumor cells with cellular proliferation rates through regulation of specific AS events, which might serve as potential therapeutic targets.

### CSFs are potentially regulated by MYC, FOX and BRD family transcription factors

Next, we investigated potential upstream transcriptional regulators of CSFs, as targeting them may have a broader effect on CSFs, with the resulting changes in EP inclusion potentially improving patient prognosis. We applied four criteria to identify high-confidence upstream transcriptional regulators of CSFs (Fig. 6A). First, as an initial filtering step, we utilized a large collection of ChIP-seq datasets across cell lines curated in the TFEA.ChIP database (Puente-Santamaria et al., 2019) and shortlisted TFs whose binding was significantly enriched within the promoter regions of CSF as compared to non-critical splicing factors (nCSFs) of EP events (first column in Fig. 6B; Methods). Next, we used the KnockTF database (Feng et al., 2020), which details transcriptome changes upon TF deletion, to calculate the enrichment of CSFs relative to nCSFs among the downregulated targets following TF deletion and retained significant hits (second column in Fig. 6B, Methods). A major limitation of KnockTF is low coverage of TFs. We therefore applied two additional computational approaches to filter the TFs shortlisted based on TFEA.ChIP.

**Figure 6.**
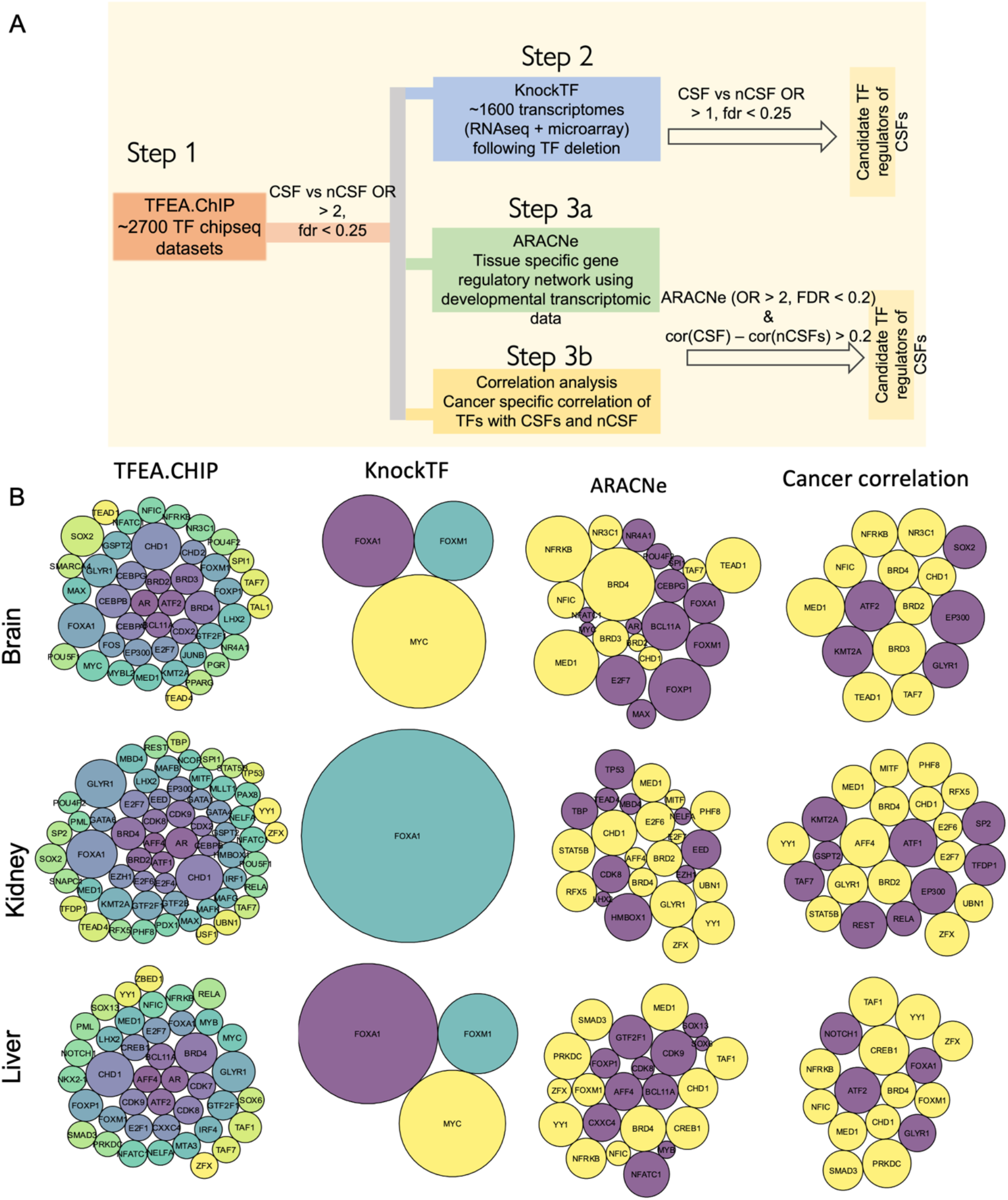
Potential TFs regulating CSFs in three organs. (A) Schematic representation of steps involved in the detection of TF regulators of CSFs. (B) Three rows correspond to three different tissues and four columns are different strategies that were used to infer TF regulators of CSFs, as labeled on the top of figures. In first three columns, bubble sizes correspond to − log10 of FDR adjusted p-values. In fourth column, bubble size corresponds to the difference between correlation coefficient of TF with median expression of CSFs and nCSFs in relevant cancer types. In first two columns, the bubbles are colored by TF names. In last two columns, yellow bubbles indicate the evidence in support of TF by both ARACNe and Cancer correlation analysis and magenta color indicates support by either one. Bubble sizes are not comparable across columns.

First, for each factor shortlisted based on TFEA.ChIP, we inferred its in-silico targets using the ARACNe software tool (Margolin et al., 2006) and selected TFs whose in-silico targets were more significantly enriched for CSFs relative to nCSFs (third column in Fig. 6B, Methods). Secondly, among the list of chip-seq filtered regulators, we identified TFs whose expression was more strongly correlated with CSFs as compared to the nCSFs in cancer transcriptomic data (fourth column in Fig. 6B, Methods). Overall, we retain in each organ, the TFs that (after the ChIP-seq based filtering) either qualified the experimental KnockTF based criterion or both of the computational filters. Collectively, these results implicate MYC, FOX (specifically FOXM1) and BRD family of TFs in the regulation of EP events through upregulation of CSFs, and may represent master regulators of broad splicing changes associated with development and cancer. Computational drug repositioning approach identified several FDA approved drugs which could potentially target these factors (Methods, Supplementary table S7).

## Discussion

The availability of transcriptomic datasets of tumors from TCGA and PCAWG consortia have facilitated the genome-wide analysis of alternative splicing changes in cancer elucidating their prognostic value (Zhang et al., 2019), genetic basis (Calabrese et al., 2020; Tian et al., 2019), and the discovery of tumor neoantigens generated by alternative splicing (Kahles et al., 2018). However, none of these studies analyzed the broader developmental context of splicing changes in cancer. Leveraging recently available temporal developmental transcriptomic data in three human organs, in this work, we have shown that the genome-wide splicing landscape of cancers significantly reverts to the early embryonic developmental stage of their tissue of origin, strongly implicating developmental splicing events in oncogenesis and tumor progression.

Similar to gene co-expression modules, inclusion of multiple exons across genes is coordinated to affect specific cellular functions during differentiation (Bland et al., 2010; Yamamoto et al., 2009), cell state transition (Warzecha et al., 2010), apoptosis (Moore et al., 2010), and hormonal induction (Hartmann et al., 2009). Our results suggest that coordinated programs of EP and EN splicing events are involved in embryonic processes, such as cellular proliferation, apoptosis, EMT, migration, and that cancers seem to misappropriate these coordinated exon inclusion events to revert to an embryonic-like state.

Further, EP and EN events ascertained based on developmental context alone are remarkably prognostic in the corresponding cancers in TCGA. For instance, the inclusion level of EP and EN events respectively predicted worse and better survival of cancer patients, underscoring the value in studying fetal development to better understand cancer mechanisms. Moreover, a negative correlation between the inclusion level of EP events and the doubling time (equivalently, positive correlation with proliferation rate) of cancer cell lines provides an independent functional validation for the role of these splicing events in mediating cellular proliferation, which is relevant to both development and cancer.

Previous research has shown that AS can affect cytoskeleton, enzymatic properties, and membrane localization of proteins (Kelemen et al., 2013). Here we observe that the molecular functions related to cytoskeleton binding and regulation of GTPase activity and cellular transport were highly enriched among EP and EN exons across all three organs we studied (Supplementary Fig. 3A). These molecular functions are central to cellular proliferation through regulation of cell cycle (Bunnell et al., 2011; Collins et al., 2012; Forth & Kapoor, 2017; Hawkins et al., 2010; McNeill et al., 2020) and cellular migration (Callan-Jones & Voituriez, 2016; Seetharaman & Etienne-Manneville, 2020; Svitkina, 2018) and, consequently, have emerged as important players in cancer progression and metastasis (Brayford et al., 2015; Buda & Pignatelli, 2011).

The analysis of protein domains enriched among EP and EN events further suggested their functional coordination in regulating diverse cellular processes such as proliferation, migration, neuronal physiology, and stress resistance. For instance, proliferation and migration of cells relies on alterations in the cytoskeleton, extracellular matrix, and cell adhesion, which are modulated by N-glycosylation of proteins like actin, cadherins and integrins (Pinho & Reis, 2015). Our observation that subunits of OST (including TUSC3 and RPN2) undergo coordinated splicing of their TRDs among EP events suggests the novel role of this splicing in the regulation of N-glycosylation during organogenesis (Fig 3C). Further TRDs in vesicle trafficking (PAQR3, PRAF2, SEC22) and mitochondria (ABCB6, ABCB8, and SDHC), WD40 in E3 ligase involved in protein degradation (DTL, FBXW9, WDR48), and NDs in GTPases for membrane signaling (ARF4, TESK1) were coordinately spliced among the brain EP exons. This suggests the functional coordination in energy metabolism and protein synthesis/processing during neuronal development is, in part, mediated via alternative splicing. In accordance, knocking down CSFs that regulate brain EP events result in the defects in the nervous system development in mice (Fig S4A), supporting the essential role of coordinately spliced EP events in organ development.

The EP and EN-mediated functional coordination is further illustrated in the case of coordinately spliced protein domains among the EN events in brain (Fig. 7B). Coordinately spliced ND in the GTPases involved in vesicle trafficking (ARL1 and RAB family genes), TRD in the endoplasmic reticulum-associated proteins involved in the ceramide/inositol synthesis (CDITP, CERS2, KDSR, etc.) and synaptic proteins involved in neuronal singalling (DAGLB, KCNN2, MCTP1, etc.) suggests the coordination in post-natal neuronal function such as setting up and firing rapid action potentials (Fig. 7B, right panel) and loss thereof in cancer (Fig. 7B left panel and Fig S6B).

**Figure 7.**
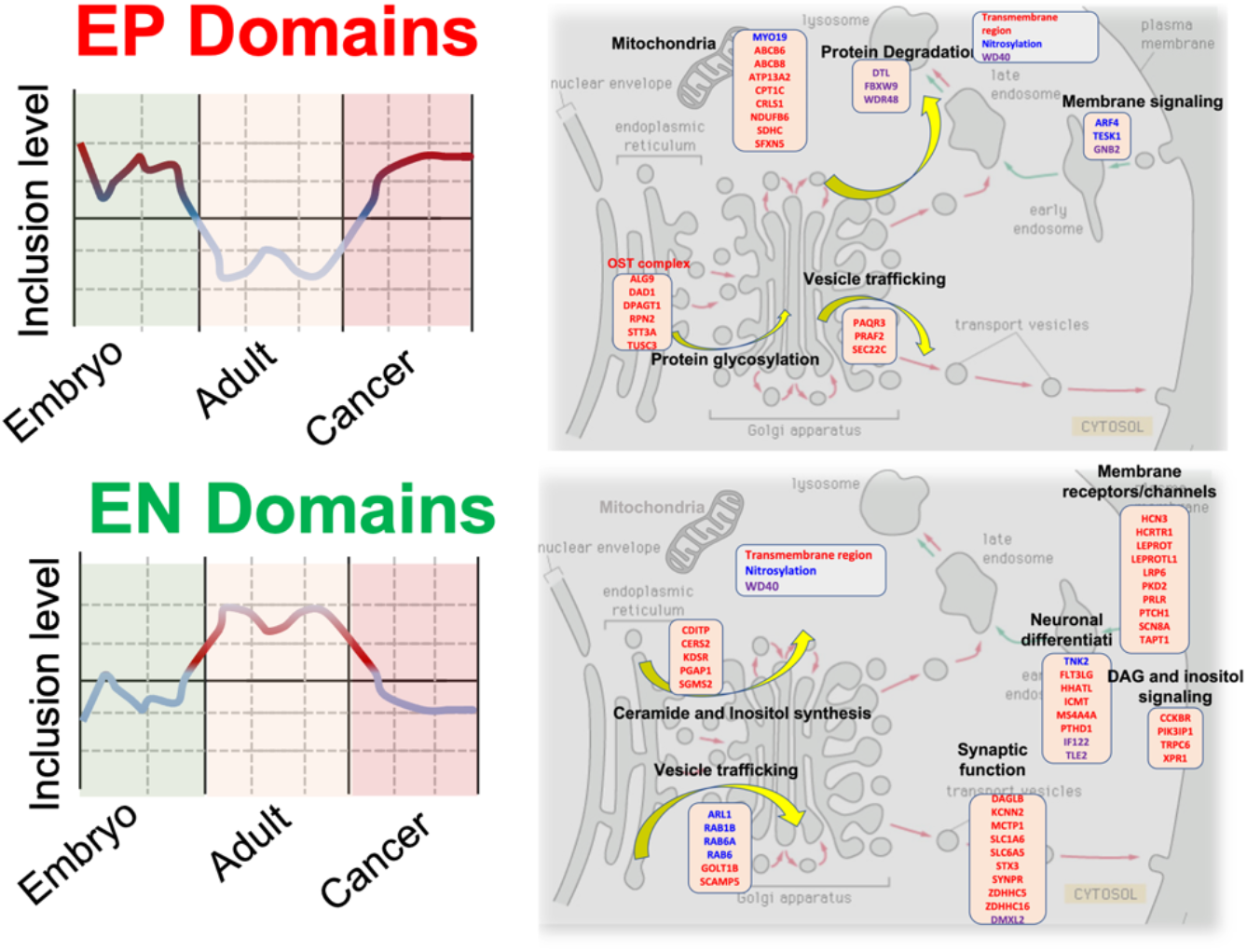
Coordinated splicing in brain development and cancer. This schematic shows the proposed role of coordinately spliced TRD, ND, and WD40 domain among EP and EN events in mediating the N-glycosylation and retrograde transport functions. (A) *Left panel,* inclusion level of EP domains in different developmental and pathological stages. *Right panel,* the cognate genes of EP events are enriched for oligosaccharyltransferase (OST) complex, vesicle trafficking, and mitochondria (transmembrane-region domain), protein degradation (WD40 domain), and membrane signaling (nitrosylation domain). These processes suggest active protein synthesis, processing including N-glycosylation, and energy metabolism during neural cell development. (B) *Left panel,* inclusion level of EN domains in different developmental and pathological stages. *Right panel,* the cognate genes of EN events are enriched for vesicle trafficking (nitrosylation domain) and neuronal function, including ceramide/inositol synthesis, synaptic function, and membrane receptors/channels/signaling (transmembrane-region domains). Most of the genes have function required for mature neural cells (e.g. neural transmission and synaptic signaling).

Moreover, for a vast majority of the EP domains, their inclusion level, which is higher in pre-natal stages, switches back to pre-natal stages (Fig 7A and S6A). This suggests that cognate genes containing these domains drive cancer progression and EP exons of these genes can be potential therapeutic anti-cancer targets.

Several single cell RNA-seq studies in the recent past have noted a general similarity in development and cancer (Couturier et al., 2020; Curry & Glasgow, 2021; Marie et al., 2020). Therefore, our results indicate that these similarities are hinged upon a much broader and coordinated reprogramming of splicing in cancer cells back to their embryonic counterparts.

Remarkably, critical splicing factors, which were inferred to regulate the inclusion of EP events based solely on the embryonic developmental data, are upregulated in cancer and confer poor prognosis to the patients. Furthermore, inactivating mutations of critical splicing factors result in the decreased level of embryonic splicing, strongly supporting their causal role in regulating embryonic splicing events. The causal role of CSFs also supported by their significantly higher, and experimentally quantified, dependency scores in the DepMap dataset in liver cancer derived cell lines. However, we did not see this trend in the brain and kidney, which may be attributed to divergent physiology and regulatory networks in cell lines, as compared to tumors in the context of the tumor microenvironment. Taken together, these observations highlight the potential of our inferred critical splicing factors to be used as therapeutic targets against cancer progression.

Further we found that CSFs in each developing organ, as well as the corresponding cancer, were likely regulated by FOX (FOXM1), MYC, and BRD family of transcriptional regulators. Regulation of splicing factors and splicing events by Myc has been previously noted (Park et al., 2019; Phillips et al., 2020). A recent report shows that MYC driven splicing factors regulate ~4000 splicing events across cancers (Urbanski, 2021). Interestingly, the FOX and the MYC family of regulators control growth, proliferation, and survival of cells in multiple contexts during embryogenesis as well as cancer (Dang, 2013; Golson & Kaestner, 2016). Our work extends the previous studies by showing regulation of splicing factors and functionally coordinated splicing events by MYC (as well as BRD and FOX family of TFs) in the developmental context, thus providing further mechanistic links between development and cancer. Our results suggest that the broad changes in the expressed isoforms of key genes driven by the upregulation of CSFs is likely a major mechanism by which these TFs exert their physiological effects. Therefore, targeting the upstream regulators of CSFs might result in broader changes in genome-wide splicing and improve the survival rates of patients. But such an approach is likely to suffer from unintended side effects. Therefore, direct targeting of EP exons, through recently developed CRISPR-based techniques (Gonatopoulos-Pournatzis et al., 2020; Thomas et al., 2020), as opposed to their upstream regulators, might result in specific lethality in the tumor cells. In the future, transcriptomic experiments following the deletion of CSFs or their upstream regulators would be necessary to establish the proposed mechanistic links and explore their therapeutic potential.

Collectively, our multi-pronged investigation not just conceptually enhances the understanding of broad functional roles and regulation of alternative splicing in the context of development and cancer, but also suggests novel putative cancer therapeutic targets. Our work also provides a framework to study the cellular mechanisms implicated in development and cancer using other molecular modalities such as miRNA and lncRNA activities, DNA methylation and histone modification profiles, alternative promoter and poly-A usage.

## Methods

### Datasets and quantification of exon inclusion

For brain, liver and kidney, uniformly processed RNA-seq data for tumors from TCGA (https://www.cancer.gov/tcga) and normal samples from GTEx (Gtex, Carithers & Moore, 2015) were downloaded from the UCSC xena browser (data version V7). The UCSCXenaTools library in R (S. Wang & Liu, 2019) was used to download transcript-level TPM values computed using Kallisto (Bray et al., 2016); the details data integration and processing can be obtained from UCSCXena browser (https://xenabrowser.net/). In total, we obtained the expression levels of 197,046 transcripts across all samples. The number of samples obtained are: brain cancer – Lower grade glioma (LGG): 523; Glioblastoma GBM: 172, normal brain – brain cerebellum: 118; brain cortex: 107, liver cancer – Liver hepatocellular carcinoma (LIHC): 369, normal liver –110, kidney cancer – Kidney renal papillary cell carcinoma (KIRP): 321; Kidney renal cell carcinoma (KIRC) 595, normal kidney – 27. For developmental data (Cardoso-Moreira et al., 2019), we obtained the raw reads from the SRA repository using the accession number E-MTAB-6814 and computed the transcript level TPM values using Kallisto (Bray et al., 2016) and the transcriptome index based on gencode version v23 annotations, the same version which was used by UCSC-XENA. The data includes multiple pre-natal and post-natal time points in each organ (Supplementary Table S1). To quantify the inclusion level of exons in each sample, we calculated the ‘percent-spliced-in’ (PSI) value for each exon, which ranges from 0-1 (i.e. from fully excluded to fully included), using SUPPA-2 (Trincado et al., 2018). Additionally, transcript-level TPMs were converted to gene-level TPMs and subsequently quantile normalized as needed for the follow-up analyses. All the scripts used for downloading and processing the RNA-seq datasets are available in at https://github.com/hannenhalli-lab/AltSplDevCancer.

### Developmental splicing events

To identify splicing events deemed to be involved in embryonic development, we adapted a previously published strategy called PEGASAS (Phillips et al., 2020). PEGASAS identifies the alternative splicing events that correlate with the activity of a specific biological pathway. In this study we identified developmental exons via a three-step process as follows.

**Step 1**: We scored the activity of each of the 332 KEGG pathways (Kanehisa et al., 2012) at each time point during development using the median of log-transformed expression of its constituent genes, resulting in a 332 x N activity matrix, where N is the number of developmental time points that were sampled for each tissue and are given in Supplementary Table S1. Clustering this activity matrix reveals two broad clusters (Supplementary Fig. S1A) – one active pre-natally and the other active post-natally.

**Step 2:** We applied an additional smoothing procedure in PCA space where our goal was to quantify each pathway’s tendency to be preferentially oriented towards a specific developmental timepoint. In 5-dimensional PC space (first 5 PCs explain ~65 % of variance), each timepoint occupies a unique coordinate based on the PC scores. In this space, similarly, each pathway corresponds to 5-dimensional vector of the pathway’s loading in each of the 5 PCs. We quantify the preferential orientation of a pathway toward a specific timepoint as cosine similarity between the loading vector and the location of the time point in the 5-dimensional space. This procedure yields a smoothed 332 x N matrix clearly segregating 332 pathways into two broad groups based on their preferential activity during pre- or post-natal stages of development (Fig. 1A). The grouped pathways were correspondingly called embryonic positive and embryonic negative pathways.

**Step 3:** Next, we used an approach similar to PEGASAS (Phillips et al., 2020) and computed the cross-sample Pearson’s correlation coefficient (PCC) between the PSI value of each exon and pathway activity score in Step 1. For each exon, we selected the significantly positively or negatively correlated KEGG pathways based on the Benjamini-Hochberg FDR threshold of 0.05. We call an exon embryonic positive (EP) if it is significantly correlated with at least 10% of the embryonic positive pathways vs. at most 5% of the embryonic negative pathways. Analogous criteria were applied to define embryonic negative (EN) exons.

This approach is superior in detecting the splicing events relevant to embryonic development of tissues compared to simply performing differential splicing between pre- and post-natal stages of development because (i) the sample size of the developmental dataset is insufficient for a robust inference, (ii) an individual exon’s inclusion can be highly variable within pre- and post-natal stages, which can confound the identification of embryonic splicing events, (iii) since this approach is anchored on robustly identified embryonic positive and negative pathways, instead of relying only on individual event’s temporal dynamics, it is likely more robust to noise.

### Cancer-specific splicing events

For each exon skipping event identified by SUPPA2, we performed a tumor-normal comparison of its PSI value to identify the splicing events which were differentially included in tumors. Owing to the transcriptomic heterogeneity across tumors, a standard differential splicing analysis, which assesses the significance of difference in the median PSI values of cancer and normal samples, will not detect exons mis-spliced in a small number of tumors, which can nevertheless be biologically significant (Kahles et al., 2018). Therefore, we selected the events which were at least 2 standard deviations away from the mean of their distribution in the corresponding GTEx normal samples in a consistent direction (i.e. increased or decreased) in at least 15% of the cancer patients. (Fig. 1A). Correspondingly, such events were termed as frequently increased or decreased in cancer.

### Comparison of cancer and developmental splicing and functional enrichment analysis

To assess if cancer recapitulates embryonic splicing events, we assessed the significance of overlap between cancer and developmental splicing events using Fisher’s exact test and adjusted the P-value using Benjamini-Hochberg’s FDR method. Functional enrichment analysis was performed using the clusterProfiler library in R and the p-values of the resulting significant terms were adjusted with Benjamini-Hochberg’s method. For plotting, the resulting GO terms were simplified based on their semantic similarity using the ‘simplify’ function from clusterProfiler in R (similarity threshold of 0.7).

### Protein domain enrichment in EP and EN exons

We downloaded the transcriptomic coordinates of all the PFAM domains mapped to the reference genome (hg38) from the prot2hg database (http://www.prot2hg.com). Since any given domain can be incorporated either fully or partially in multiple transcripts, the downloaded file was preprocessed to remove redundancy of genomic coordinates resulting from the same domain mapping to multiple transcripts by using bedtools merge. We then intersected the preprocessed genomic coordinates of protein domains to the unique and non-overlapping set of EP and EN exons as well as the rest of the alternatively spliced skip exons (called background exons) using bedtools intersect in each tissue. To identify the domains enriched in EP and EN events, we computed the frequency of occurrence of each domain in EP, EN, and background exons and performed a Fishers’ test of enrichment in each tissue. The resulting p-values from the Fishers’ test were corrected for multiple testing by using Benjamini-Hochberg’s method and the domains with an odds ratio > 1 and corrected p-value < 0.1 were considered enriched among EP or EN exons in each tissue.

### Survival analysis

We used clinical data from TCGA to model the overall survival of cancer patients using the inclusion level (PSI value) of each exon (Ei) as a predictor variable and age as a covariate in the cox regression:

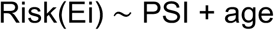

We used the R library “survival” for this analysis and the resulting p-values were adjusted for multiple testing using Benjamini-Hochberg’s method. The distribution of the resulting hazard ratios was compared between embryonic positive, negative and the rest of the splicing events.

### Model for regulation of embryonic splicing

To dissect the potential regulators of embryonic splicing events, we built upon a commonly used notion that differential expression of splicing factors could lead to the differential splicing of the exons (Z. Wang & Burge, 2008). For this, we identified 442 proteins which have the term ‘splicing’ in their GO definition from the Amigo database (Carbon et al., 2009). We then used a partial least square regression (PLSR) analysis to model the inclusion of EP events using the gene expression of splicing factors in the developmental data. PLSR outperforms multiple linear regression when dealing with multicollinearity among the predictor variables or when the predictor matrix is non-singular (Bjørn Helge & MevikRon Wehrens, 2007).

For gene expression matrix X of 442 features (SFs) across N developmental timepoints (n x 442) and response matrix Y of median EP splicing across n timepoints (n x 1), the PLSR transforms X and Y as per the following relations:

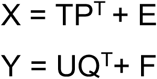

Where T and U are the N x r matrices of the extracted latent vectors and P (p x r) and Q (1 x r) are the loadings of X and Y. E (n x p) and F (1 x p) are the residuals. In the PLSR algorithm, T and U are constrained to have a maximum covariance as per following relation:

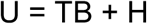

Where B (r x r) is a diagonal matrix of regression coefficients and H is a matrix of residuals. Splicing factors with positive regression coefficients and a significant p-value (p < 0.05 after FDR correction) were considered critical regulators of EP events (CSF) PLSR was implemented using ‘pls’ package in R (Mevik et al., 2016).

### Mutation analysis of splicing factors

To assess the causal role of CSFs in the regulation of EP events, we obtained level 2 mutation data from TCGA cohorts of brain, liver, and kidney cancers (https://portal.gdc.cancer.gov/) and identified the tumors which had nonsense or truncating mutations for these factors. For each mutated factor in each cancer type, we compared the median inclusion level of EP events in the mutant samples against the background set of samples that were not mutated for any of the splicing factors. Thus, the factors were classified into ‘increased’ or ‘decreased’ categories depending upon at least 5% increase or decrease in the median EP inclusion level. To account for the potential confounding effect of the differential expression of splicing factors between samples, we identified, for each mutant sample, a set of 10 non-mutant samples with similar splicing factor expression. Specifically, for each mutant sample, we identified 10 non-mutant samples with the shortest Euclidean distance to the mutant sample in terms of the gene expression of all splicing factors. For robustness, we discarded the splicing factors for which the background set of patients had a high variability (Std. Dev. > 0.1) in the median EP splicing across the 10 samples (Supplementary Fig. S4B).

### Transcriptional regulators of splicing factors

To identify the potential transcriptional regulators of critical splicing factors (Fig. 6A), in each organ independently, we divided the splicing factors into two classes: namely, a foreground set comprising of the top 100 critical splicing factors, and a background set comprising the remaining splicing factors (nCSFs). To assess whether a TF was more likely to regulate CSFs compared to nCSFs, we used four complementary approaches (Fig. 6A). In the first step, we used the TFEA.ChIP library in R, which uses publicly available genome-wide binding datasets from ChIP-seq experiments (Puente-Santamaria et al., 2019). TFEA.ChIP used a Fisher’s test to assess if a specific TF’s binding is significantly enriched in the promoter regions of the CSFs relative to nCSFs (step 1 in Fig. 6A). TFs with an odds ratio > 2 and an FDR of 0.05 were considered putative regulators of CSFs. This first step was used as a strict filter for a TF to be further considered. To validate the ChIP-seq-based findings with the gene knockout/knockdown studies, we used the knockTF database, which is a compendium of publicly available genome-wide transcriptional profiling following the deletion of TFs across multiple cell lines (step 2 in Fig. 6A). In this step, we obtained all the genes which were marked as downregulated based on a robust statistical analysis in the knockTF database (Feng et al., 2020) following the deletion of a transcription factor and again assessed if CSFs were enriched as compared to nCSFs among the downregulated targets using a Fisher’s test. TFs with an FDR of < 0.25 and a positive odds ratio in any of the cell lines were considered putative experimentally-derived regulators of CSFs. Furthermore, because knockTF has a poor coverage of TFs, we did not use this as a strict filter and instead used two additional computational approaches to infer the potential TFs: (i) We built a gene regulatory network for TFs shortlisted by ChIP-seq using the developmental time course data for relevant tissues and the ARACNe software (Margolin et al., 2006) and assessed if the CSFs were enriched relative to nCSFs among the in silico derived targets of each TF using a Fisher’s test (step 3a in Fig. 6A). TFs with an odds ratio > 2 and a FDR < 0.2 were considered potential in silico derived regulators of CSFs. (ii) In parallel, we assessed the correlation of ChIP-seq shortlisted factors with CSFs and nCSFs in relevant cancer types (step 3b in Fig. 6A). The factors, with a correlation difference > 0.2 between CSFs and nCSFs were considered putative regulators. The ChIP-seq shortlisted factors, which either passed the knockTF test OR passed both computational tests, were proposed as regulators of CSFs. In all the applicable cases, p-values were adjusted for multiple comparisons using the Benjamini-Hochberg procedure in R.

### Computation drug repositioning

We performed virtual screening process using Autodock Vina (Trott & Olson, 2009). The 3D structure of the FOXM1 and MYC proteins were downloaded from the RCSB-PDB (Berman et al., 2000) and further refinement was done using Open Babel software (O’Boyle et al., 2011) and the drug library was created by downloading the molecules from the ZINC database (Irwin & Shoichet, 2005) where we considered only those drug molecules which are either FDA approved or are under clinical trials. The ligand and the receptor files were prepared in the ‘pdbqt’ file format, and the center and the grid size of the receptor molecules was computed using UCSF Chimera software (Pettersen et al., 2004). Based on the affinity score and RMSD value, we proposed some potential drug molecules.

### CNV analysis

For each cancer type we obtained the level 4 CNV data from TCGA, which contained sample specific information about the CNV profile of each gene (1 being CNV amplification, 0 being no CNVs, −1 being CNV deletion). To assess the CNVs of CSFs in each cancer type, we divided all samples into three quartiles based on the gene expression of each CSF. For each group of samples obtained in this way, we calculated the average CNV value for each CSF and compared these values for all CSFs between the quartiles using a Wilcoxon test.

### Single cell validation

For single cell validation of prioritized transcription and splicing factors, we obtained GBM single-cell SMART-seq datasets from 20 adult GBM tumors (Neftel et al., 2019) from the Broad Institute Single Cell Portal (https://singlecell.broadinstitute.org/single_cell; Accession: SCP393). We also obtained normal brain single-cell SMART-seq and RNA-seq data and the annotations of cells from multiple cortical areas of the human brain from the Allen Brain atlas (2019 SMART-seq release, https://portal.brain-map.org/atlases-and-data/rnaseq). Oligodendrocytes, astrocytes, and oligodendrocyte progenitor cells were used as a normal reference to compute log-fold changes between malignant and normal cells. For liver cancer, LIHC single-cell RNA-seq data is 10X data sourced from a previous study (Ma et al., 2019) and the read count matrices and annotations were downloaded from the GEO database (GSE125449). For healthy liver, read count matrices were obtained from the HumanLiver package (https://github.com/BaderLab/HumanLiver, MacParland et al., 2018). Hepatocyte clusters (Hep 1 - 6) and cholangiocytes were used as a normal reference to compute log-fold changes between malignant and normal cells. The activity of CSFs as a single-cell level was scored as a gene set using AUCell (Aibar et al., 2017), and the resulting activity scores were z-scored across all cells separately in each tissue.

## Supplementary Figures

**Figure S1.**
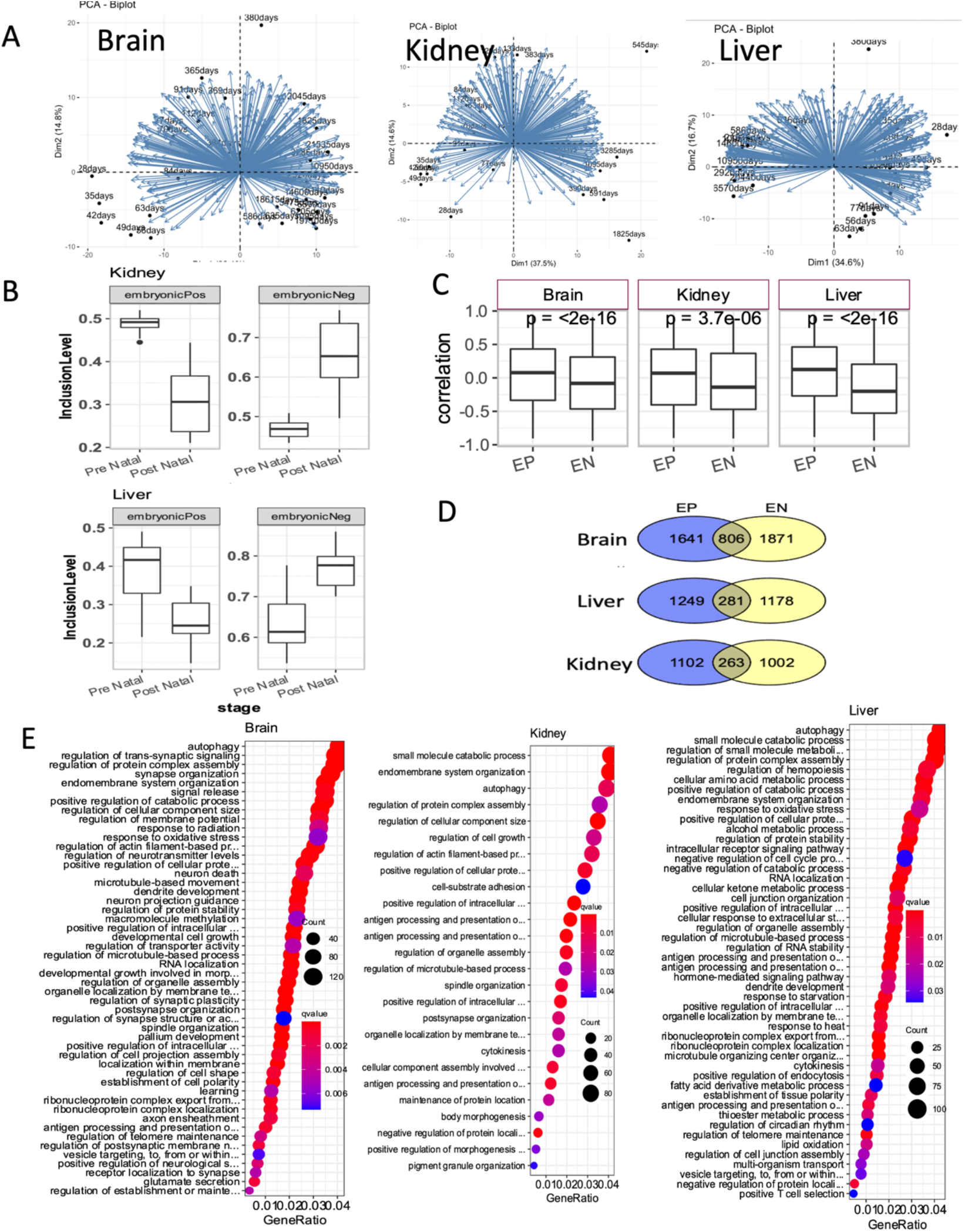
(A) Biplot from the principal component analysis of the KEGG pathway scores during the developmental timeline of three human tissues. (B) Boxplots showing the differential inclusion of embryonic positive (EP) and embryonic negative (EN) events during pre- and post-natal stages of development for kidney and liver (related to figure 1E). (C) Overlap of the cognate genes of EP and EN events in three tissues, emphasizing that same gene might contain EP and EN events. (D) Dotplots for the GO terms (biological processes) enrichment among the cognate genes of EP and EN events in three tissues.

**Figure S2.**
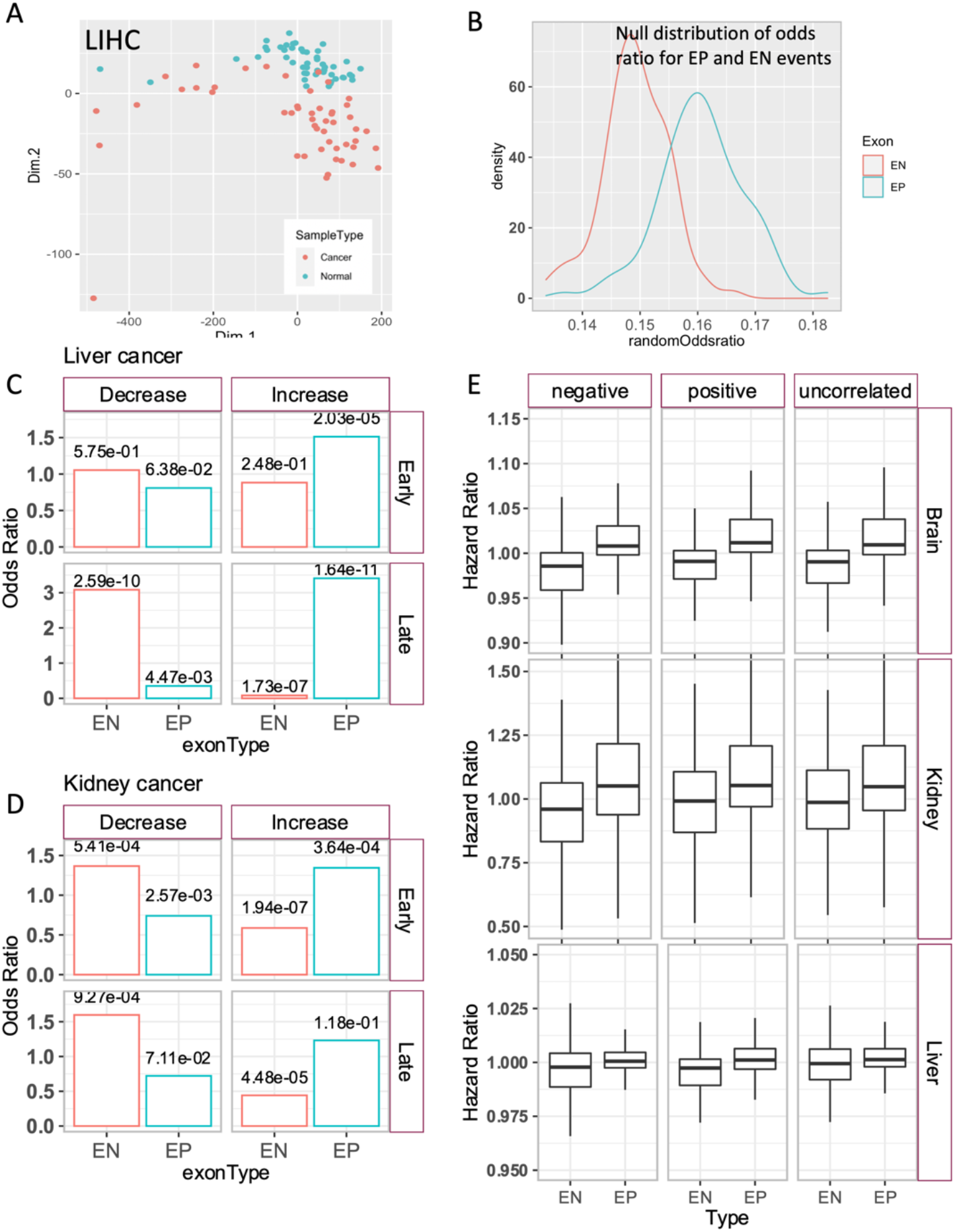
(A) Biplot from the principal component analysis of the inclusion level (PSI value) of exon skip events in liver cancer. (B) Null distribution for the odds ratio of enrichment of randomly sampled exons among the events frequently increased and decreased in cancer. These random exons were samples to have low inclusion (PSI < 0.30) and high inclusion (PSI > 0.70) for EP and EN events, respectively, in Gtex tissues, ensuring that enrichment of EP and EN events in cancer is not resulting from their baseline inclusion level in adult tissues. (C,D) Odds ratio for the enrichment of EP and EN events among frequently increased and decreased events in the early and late stage cancers, showing the greater reversion to embryonic splicing in advanced stage cancers. (E) Hazard ratio for EP and EN exons separated into three subclasses; namely negative, positive, and uncorrelated depending on their correlation with the cognate gene, showing that HRs of EP and EN exons is is independent of the expression of their cognate gene.

**Figure S3.**
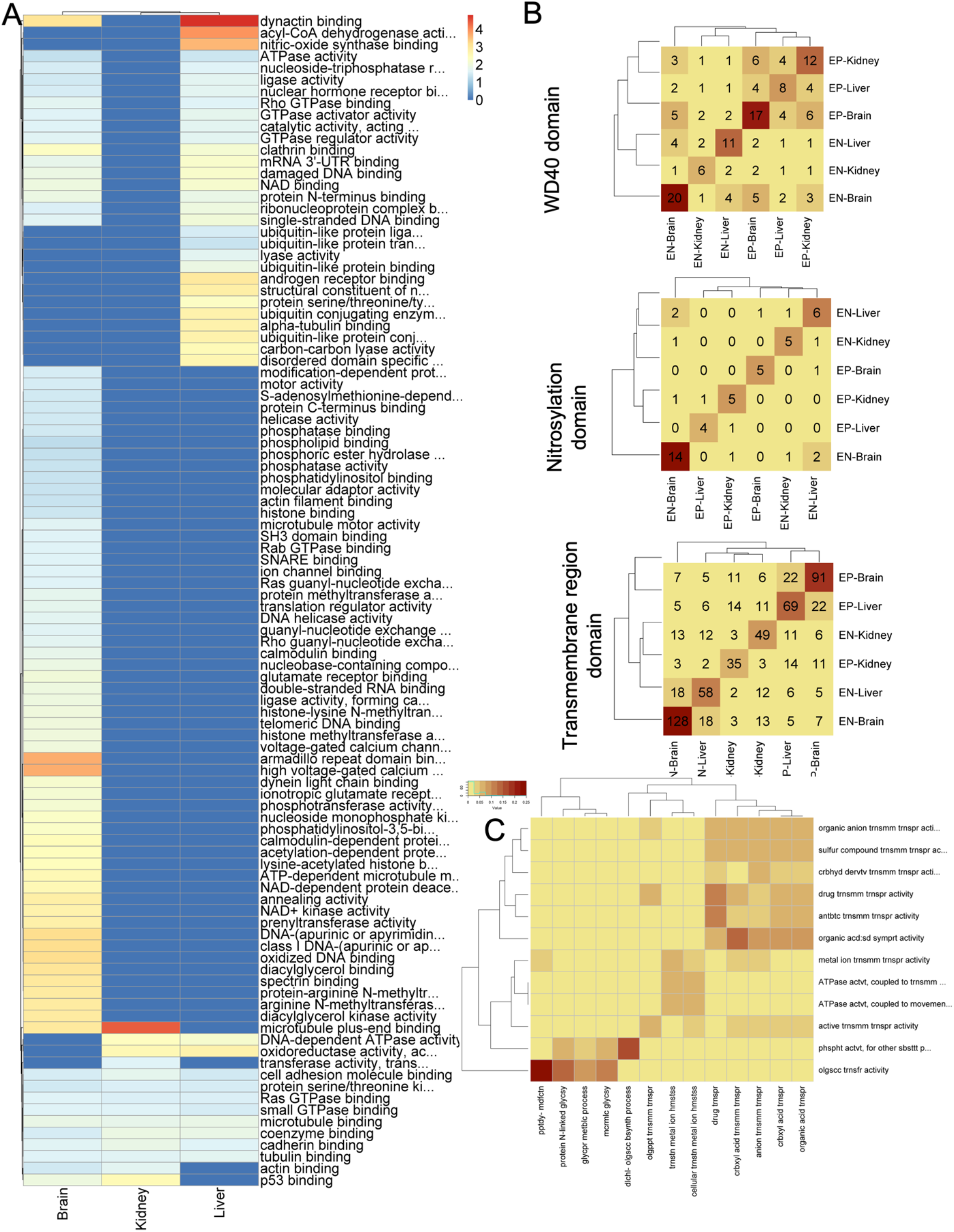
Heatmap for the GO term (molecular function) enrichment analysis of cognate genes of EP and EN events in three tissues. The colors in heat cell correspond to −log10 of FDR corrected p-value of enrichment. (B) Heatmap for the overlap of gene sets enriched for WD40, nitrosylation and transmembrane-region domains across EP and EN events in three tissues. For each domain (i.e three plots), the numbers along the diagonal indicate the number of genes having that specific domain and off-diagonal entries show the overlap between the cognate genes of EP/EN events across tissues. This plot emphasizes that for a given domain, the observed enrichment of these three domains (Fig 3A) is driven by different set of genes. (C) Correspondence between molecular functions and biological processes for transmembrane domains in brain EP events.

**Figure S4.**
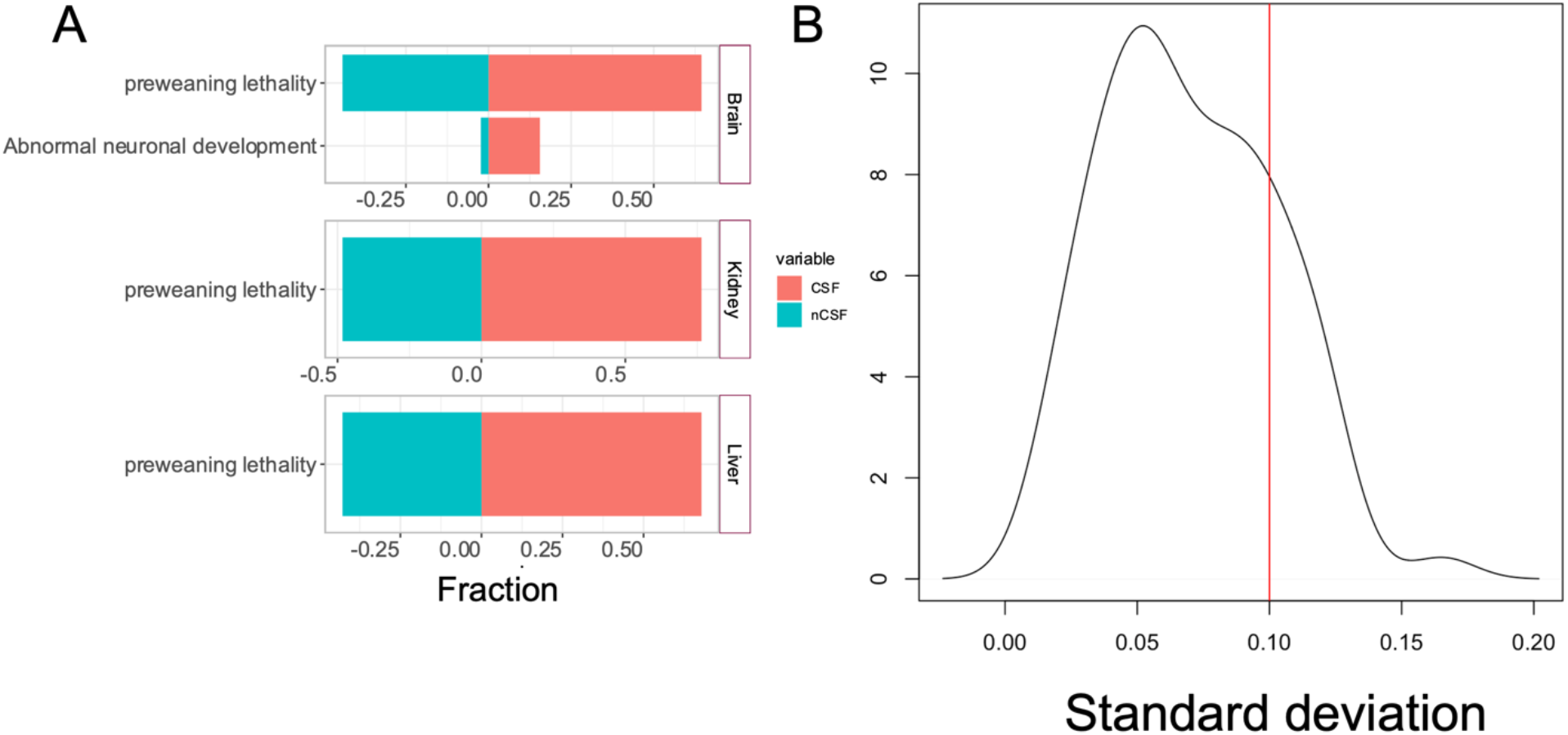
(A) Phenotypic consequences of deletion/knockdown of CSFs and nCSFs orthologs in mice. Barplots show the fraction of CSFs (red) and nCSFs (blue) that results in phenotypic defects (shown next to bar plots) in mice. The deletion of brain CSFs results in abnormal nervous development. CSFs from all three tissues are much more likely to result in pre-weaning lethality as compared to CSFs. Phenotypes shown here had an enrichment of CSFs as compared to nCSFs at pvalue threshold of 0.1 and FDR of 0.30. For this analysis, we complied the list of phenotypes associated with genetic deletion screens in mice from mouse genome informatics database (http://www.informatics.jax.org/). We manually curated the phenotypes which resulted in lethality during pre-natal stage or in developmental abnormalities in brain, kidney, and liver. The complete set of phenotypes and their associated genes is given in Supplementary Table S6. (B) For each CFS, we identified samples with inactivating mutations, and for each such sample, we identified 10 wt smaples with similar expression of the CSF, computed median PSI across EP events in each sample and estimated the standard deviation across the 10 control wt samples. The figure shows the density plot of standard deviations across CSFs. To ensure the homogeneity of control samples in terms of their median EP splicing, we discarded all CSFs for which the control samples had a standard deviation of > 0.1.

**Figure S5.**
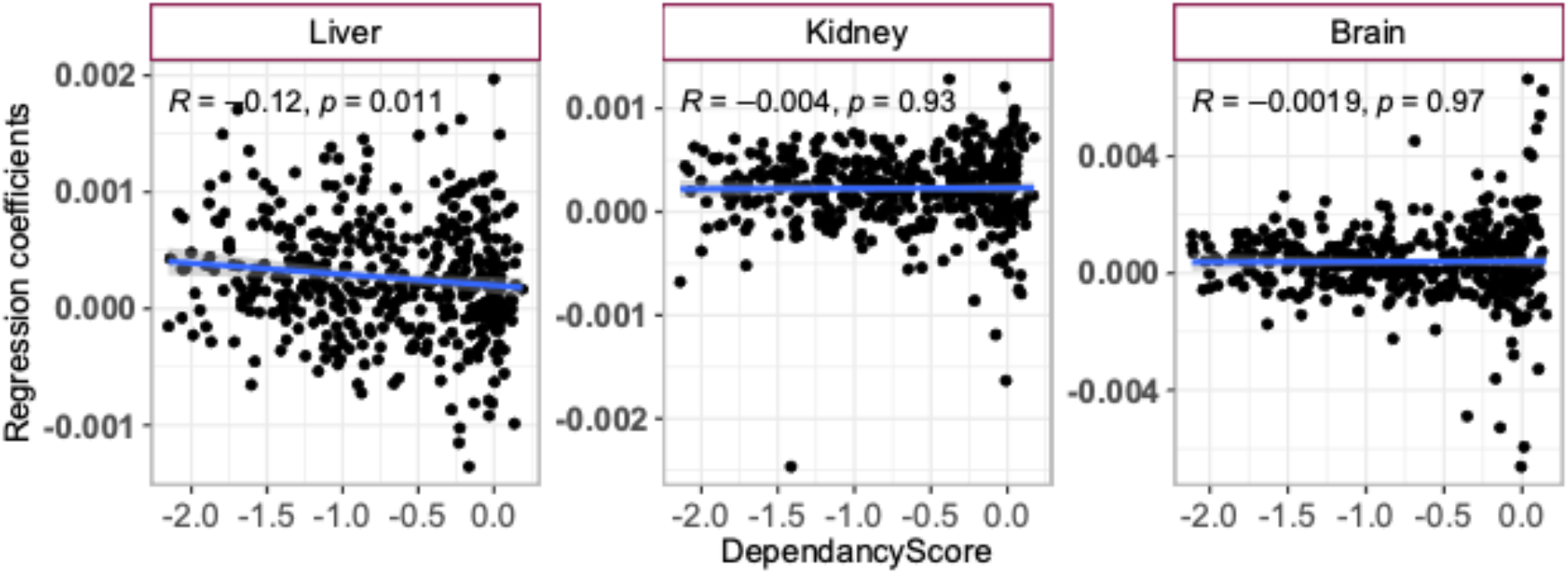
Scatter plot for the correlation of regression coefficients of CSFs with the mean dependency score of CCLE cancer cell lines from DepMap project. Each dot is a CSF.

**Figure S6:**
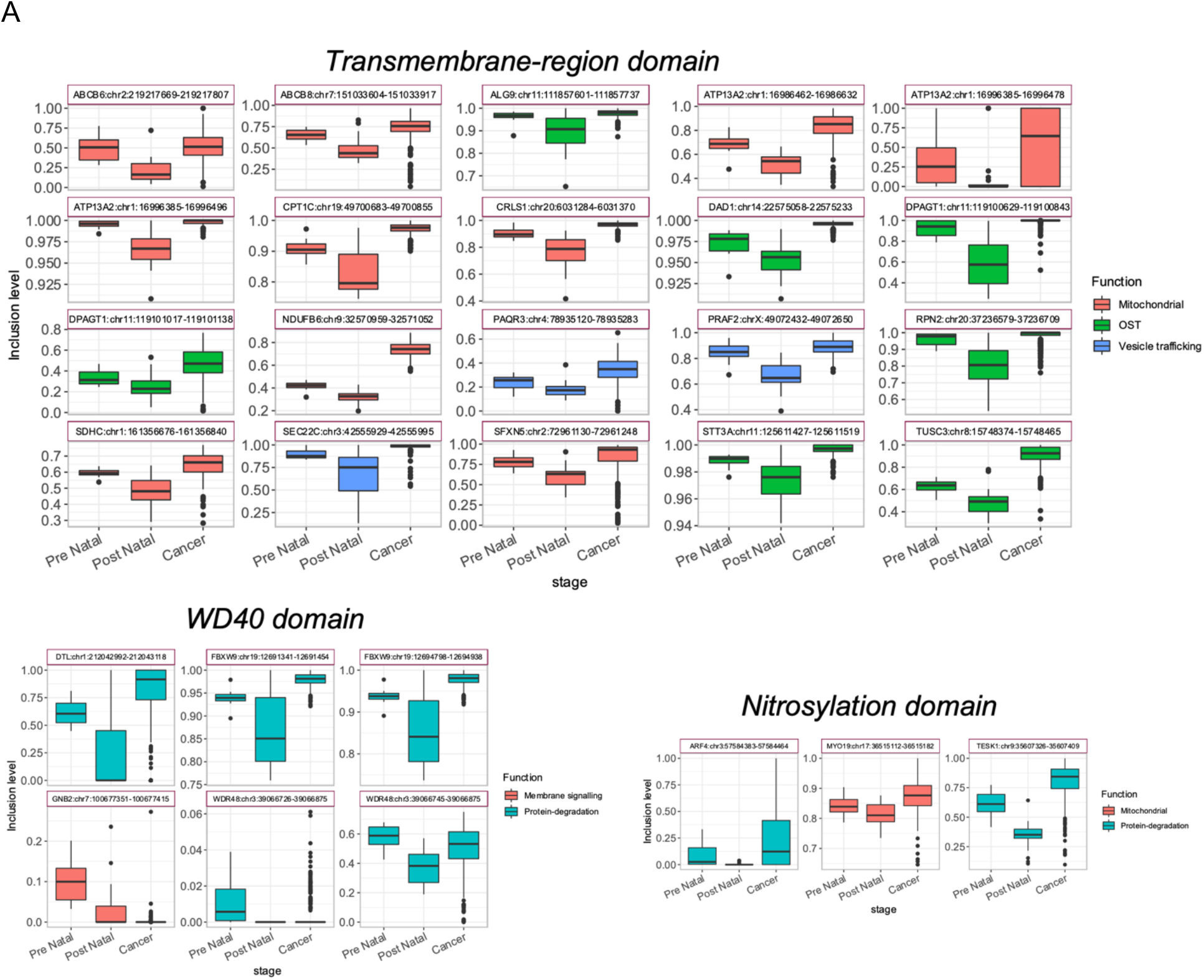
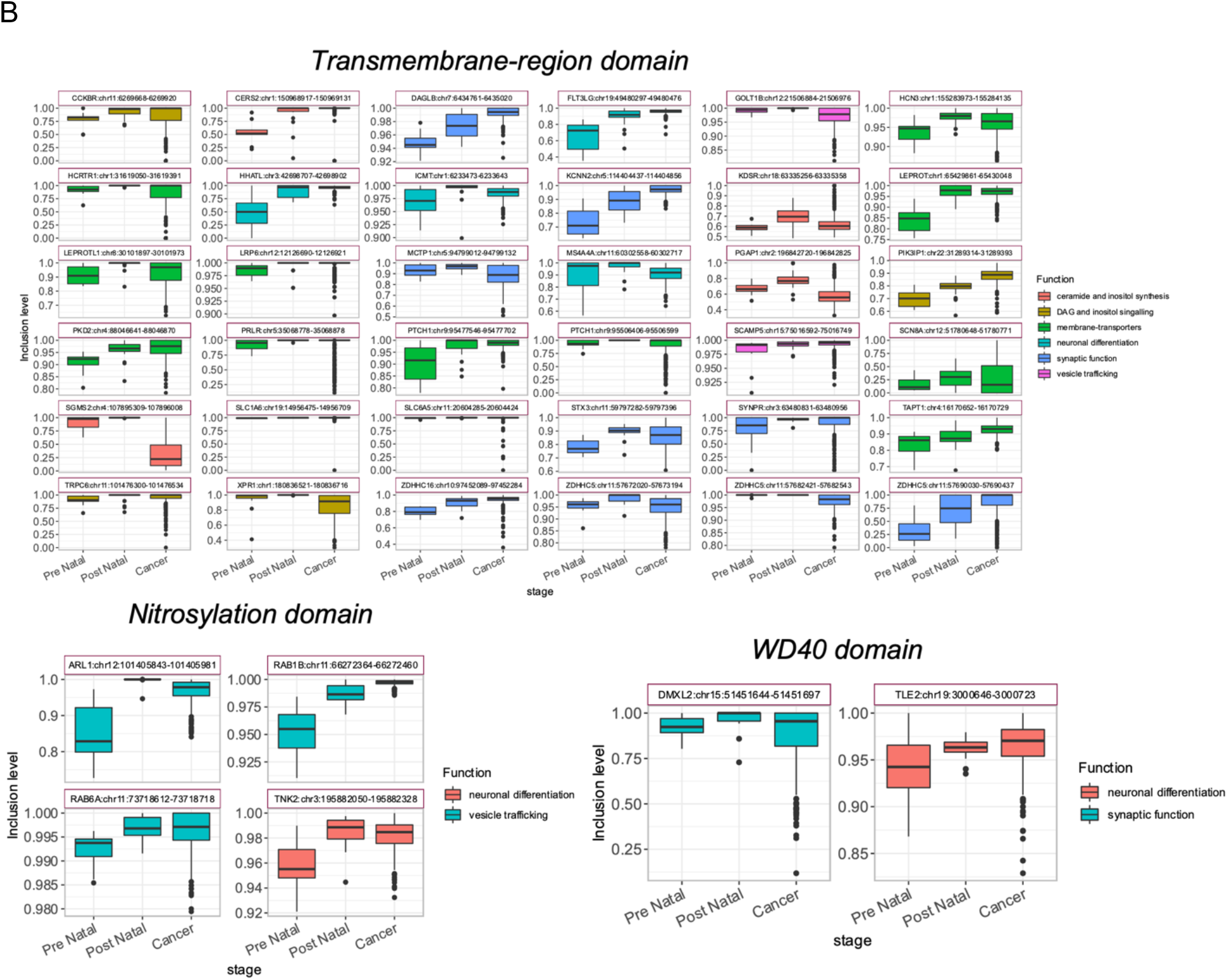
Inclusion level of functionally coordinated EP (A) and EN (B) exons containing TRD, WD40 and nitrosylation domain in pre-natal, post-natal, and cancer samples of brain. Text inside the stripes of boxplots are gene names and exon coordinates and color represents biological functions performed by the genes. A detailed description is provided in the Discussion section.

